# Transcriptome profiling implicates a role for Wnt signaling during epileptogenesis in a mouse model of temporal lobe epilepsy

**DOI:** 10.1101/2022.07.22.501153

**Authors:** Muriel D Mardones, Kunal Gupta

## Abstract

Mesial Temporal lobe epilepsy (mTLE) is a life-threatening condition characterized by recurrent epileptic seizures initiating in the hippocampus. mTLE can develop after exposure to risk factors such as seizure, trauma, and infection. Within the latent period between exposure and onset of recurrent seizures, pathological remodeling events occur which are believed to contribute to epileptogenesis. The molecular mechanisms responsible for epileptogenesis in the seizure network are currently unclear. We used the mouse intrahippocampal kainate model of mTLE to investigate transcriptional dysregulation in the ipsilateral-injected epileptogenic zone (EZ), and contralateral peri-ictal zone (PIZ) in the dentate gyrus (DG) of the hippocampus during the first 14-days after induction of status epilepticus (SE). DG were micro-dissected 3, 7 and 14-days after SE for high-throughput RNA-sequencing. In the EZ, dynamic transcriptional dysregulation was evident over 2-weeks with early expression of genes representing cell signaling, migration and proliferation. In the PIZ, gene dysregulation was most prominent at 3-days in similar domains. Inflammatory gene groups were also prominent over the 2-week epileptogenic period in the EZ and PIZ. We uncovered that the Wnt signaling pathway was dysregulated in the EZ and PIZ at 3-days and we validated these changes via immunohistochemistry. This suggests that critical gene changes occur early after neurological insult and that canonical Wnt signaling may play a role within this latent period. These findings offer new insights into gene expression changes that occur in the hippocampal DG early after SE and may help to identify novel therapeutic targets that could prevent epileptogenesis.

**Significance statement:** Mesial temporal lobe epilepsy (mTLE) is a severe life-threatening condition that is often medically refractory. While risk factors for the delayed development of mTLE are well-known, there are currently no therapeutic interventions that prevent epileptogenesis. Knowledge of the gene dysregulation events that occur during the latent period between exposure and epilepsy is critical to understanding epileptogenesis and developing new therapies. We utilized a mouse model of adult focal mTLE, the most common form of adult clinical epilepsy, and investigated transcriptional changes in the dentate gyrus during the first 2-weeks after status epilepticus. These data provide new insights into specific gene changes and pathways within different regions of the seizure network that could be targeted to prevent the development of epilepsy.

## Introduction

Epilepsy is a common neurological disorder that afflicts 1-2% of the US population; the most common form of adult focal epilepsy is mesial temporal lobe epilepsy (mTLE), which is medically refractory in 40% of cases (Kwan, 2000; Kwan, 2004; Kobau, 2008; Picot, 2008). In many cases, an inciting event or risk factor (e.g. febrile seizure, trauma, infection) can be identified prior to the onset of clinical epilepsy (Frey, 2003; Myint, 2006; French, 2012; Ramantani, 2017; Chen, 2018a). The process of epileptogenesis, which occurs in this latent period, is characterized by pathological remodeling of the hippocampus, specifically the dentate gyrus. Critically, rodent models of epilepsy recapitulate important aspects of human mTLE, such as mesial temporal sclerosis, hippocampal dentate granule cell dispersion, dentate granule cell mossy fiber sprouting and the loss of hippocampal hilar and pyramidal neurons (Parent, 1997; Kralic, 2005; Kron, 2010; Cho, 2015). Furthermore, induction of a single episode of status epilepticus (SE) by a variety of established methods in these rodent models typically results in acquisition of epilepsy within two weeks (Twele, 2016a; Twele, 2016b; Twele, 2017), affording a unique opportunity to investigate the earliest stages of epileptogenesis *in vivo.*

It is increasingly well recognized that the epileptic brain develops established seizure propagation pathways that predisposes to recurrent seizures (Wendling, 2010; Nadler, 2014; Sheybani, 2018), and persistence of these epileptic circuits after surgery may be responsible for delayed surgical failures (McIntosh, 2004; Barba, 2016). The molecular mechanisms responsible for epileptogenesis and seizure circuit formation represent unique therapeutic targets that might prevent epileptogenesis after an injury event and may even improve the efficacy of clinical interventions. As a result, prior studies have sought to investigate transcriptomic and proteomic changes in human and animal models of epilepsy (Li, 2010; Theilhaber, 2013; Hansen, 2014; Dixit, 2016; Dingledine, 2017; Canto, 2021); several studies have indeed focused on the dentate gyrus as one of the key sites in mTLE epileptogenesis (Li, 2010; Dingledine, 2017; Canto, 2021). These studies and others have contributed significantly to our understanding of epilepsy-specific gene expression changes, implicating specific molecular pathways including MAPK, NMDA-receptor-mediated excitotoxicity, CaMK, mTor and Wnt (Theilhaber, 2013; Hansen, 2014; Canto, 2021).

There is growing evidence that the Wnt pathway may provide some of the key early signaling events during epileptogenesis (Gupta, 2019; Sun, 2020; Alqurashi, 2021; Jean, 2022). Wnt signaling encompasses a family of 19 secreted Wnt protein ligands, which bind a family of 10 membrane frizzled receptors and co-receptors; these signal to downstream canonical beta-catenin-dependent and non-canonical planar cell polarity and calcium pathways (Oliva, 2013). Under physiological conditions, Wnt pathway signaling has been implicated in neurogenesis and dendrite formation in the adult rodent hippocampus (Lie, 2005; Gao, 2007; Choe, 2012; Mardones, 2016; Arredondo, 2020b). More recently, the canonical Wnt pathway has also been implicated under pathophysiological conditions during epileptogenesis, affecting pro-epileptogenic neuronal remodeling and neurogenesis in the hippocampal dentate gyrus (Fasen, 2002; Qu, 2017; Gupta, 2019).

In this study, we used the adult mouse intrahippocampal kainate model of focal mesial temporal lobe epilepsy and performed whole transcriptome profiling of the hippocampal dentate gyrus. Our use of this model provides a unique opportunity to examine transcriptional dysregulation in varying regions of the seizure network. It has been demonstrated that the contralateral dentate gyrus also undergoes pathological remodeling in this model, and it is well-recognized that the contralateral hippocampus is an independent source of epileptic seizures (Häussler, 2012; Gupta, 2019). We therefore investigated both the ipsilateral injected dentate gyrus, representing the epileptogenic zone and the contralateral dentate gyrus, representing the peri-ictal zone, recognizing the role of both parts of the seizure network in maintaining epilepsy (Wendling, 2010; Nadler, 2014; Sheybani, 2018; Gupta, 2019). We selected time-points during epileptogenesis, 3-days, 7-days and 14-days after SE, which fall within the latent period between SE and the onset of spontaneous recurrent seizures (Twele, 2016a; Dingledine, 2017; Canto, 2021). The data presented herein identify distinct patterns of gene expression dysregulation within the seizure network, which are dynamic over the early epileptogenic period: In the epileptogenic zone, we identify early enrichment of cell signaling and delayed inflammatory processes, and in the peri-ictal zone we identify a brief early enrichment of cell signaling and inflammatory processes. We further observe that the canonical Wnt pathway is dysregulated during early epileptogenesis in the epileptogenic zone and the peri-ictal zone. These findings provide clues as to the key dysregulatory events that may contribute to epileptogenesis and potentially identify novel therapeutic targets that might be modulated to prevent the development of epilepsy.

## Methods

### Animal husbandry

Experiments were performed using male and female adult wild-type C57Bl6/J mice (Jax #000664) of 8-10 weeks of age. Mice were housed in group cages with a maximum of 5 animals per cage with free access to food and water *ad libitum,* according to local IACUC guidelines. Experimental surgeries were performed at ZT3-5 Zeitgeber time. All animal procedures were performed in accordance with the [Author University] animal care committee’s regulations.

### Intrahippocampal kainate induced status epilepticus

Mice were induced and maintained under anesthesia with isoflurane by spontaneous respiration. The head was secured with ear bars in stereotactic apparatus (Stoelting), the head was shaved and the scalp sterilized with alternating ethanol and betadine. A single midline incision was made, stereotactic coordinates were obtained relative to bregma. A single burr hole was placed at X +1.8, Y −2.1, the debris was cleared with sterile saline injection. The injection needle (Hamilton) was slowly advanced to Z −1.9 from the skull. 85nl of sterile normal saline or kainate (Millipore Sigma, 18.7nM in normal saline) were delivered over 1-minute using the Quintessential Stereotactic Injector (Stoelting); the needle was retained in position for 2 minutes to prevent reflux prior to slow removal. The skin was closed using dermal glue (VetBond 3M) and the animal recovered in a warmed chamber. Seizures were scored for 2-hours after injection by modified Racine scale (references); stages 1 and 2 – freezing, head nodding, mastication, stage 3 – forelimb clonus, stage 4 – rearing, stage 5 – rearing and falling, stage 6 – “popcorn”-type seizure. Induction of status epilepticus (SE) is well-described after intrahippocampal kainate injection, therefore, seizure induction was confirmed behaviorally without the use of EEG as described previously (Kralic, 2005; Heinrich, 2006; Overstreet-Wadiche, 2006; Kiasalari, 2013; Qu, 2017; Lee, 2018). Only mice that underwent at least one observed Racine 3-6 seizure were included in the epilepsy group; all animals that received kainate incurred a Racine 3-6 seizure within the 2-hour observation period, no saline-injected mice demonstrated behavioral seizure activity. N=4 per group (2 male, 2 female). Mice were administered subcutaneous carprofen (5mg/kg) and food saturated with pediatric Tylenol (3.2mg/ml) for 24-hours after surgery and soft-food daily until tissue extraction.

### RNA extraction and library preparation

Brains were extracted 3-days, 7-days and 14-days after induction of SE/control saline injection. The dentate gyri were anatomically microdissected in an RNAse-free environment. The ipsilateral and contralateral dentate gyri were isolated separately and hemisected into dorsal and ventral parts; the dorsal dentate gyri were used for transcriptomic analysis. Dorsal ipsilateral dentate gyrus tissue represented the epileptogenic zone (EZ), synonymous with the ictal zone, and the dorsal contralateral dentate gyrus tissue represented the peri-ictal zone (PIZ). Tissues were preserved in RNALater (AM7020, Invitrogen) for 24hrs at 4°C, mechanically macerated (Kimble-Chase) and stored at −80°C until all tissues were collected. RNA-extraction was performed for all samples simultaneously utilizing high-throughput automation. Samples were prepared for extraction using the standard protocol for lipid-rich samples described in the QIAsymphony RNA Handbook (November 2020 edition). Tissue disruption and homogenization was carried out using a Qiagen Tissuelyser II. RNA extraction was performed using a QIAsymphony SP (protocol: RNA_CT_800_V7, kit: QIAsymphony RNA Kit (192), catalog no. 931636). In brief, RNA was separated from lysates through binding to magnetic particles, DNA was removed by RNase-free DNase treatment, RNA was eluted into 100 μL of RNase-free water and analyzed using an Agilent 4200 TapeStation. All samples received RNA integrity scores over 7.5. Poly-A selected libraries were prepared with the Illumina TruSeq Stranded mRNA Library Preparation Kit protocol (catalog no. 20020595). The libraries were analyzed by Agilent 4200 TapeStation, and pooled. The pooled libraries were loaded on a NextSeq 500 High Output v2.5 run with a 75-cycle sequencing module (catalog no. 20024906) to generate paired-end reads. The demultiplexing of the reads was performed with bcl2fastq version 2.20.0.

### RNA-Sequencing and bioinformatic analysis

48 poly-A selected RNA-seq libraries were sequenced to an average depth of 11.2 million read pairs. On average 9.8 million read pairs were mapped as properly paired reads (88%) per sample. Of these 9.8 million read pairs, 8.5 million could be assigned to known features (87%). NextSeq reads were trimmed using fastp (version 0.20.1) with parameters “-l 17 --detect_adapter_for_pe – g -p” (Chen, 2018b). The resulting reads were mapped against GRCm39 using HISAT2 (version 2.2.1) with parameters “--rna-strandness F” (Kim, 2015). HISAT uses Bowtie2, which is based on the Burrows-Wheeler transform algorithm, for sequence alignment and allows for mapping across exon junctions (Langmead, 2012). Read counts for each gene were created using featureCounts from the Subread package (version 1.6.4) with the parameters “-O -M --primary -p --largestOverlap -s 2 -B” and Gencode M28 as the annotation (Liao, 2014; Frankish, 2021). Differential expression analysis was performed using the DESeq2 package (version 1.32.0) in R/Bioconductor (R version 4.1.1) (Love, 2014). Functional analysis was performed using The Database for Annotation, Visualization and Integrated Discovery (DAVID) version2022q1 (Huang da, 2009; Sherman, 2022). Gene lists for each timepoint were selected using fold change > 1.5 and FDR < 0.01, then analyzed through DAVID web services using custom perl scripts. GO terms (GOTERM_XX_FAT), Kyoto Encyclopedia of Genes and Genomes (KEGG)-pathways, and BioCarta Pathways were used for the functional enrichment analysis (Ashburner, 2000; Kanehisa, 2000; 2001; Kanehisa, 2019; 2021; Kanehisa, 2021). Genes with FDR<0.01 and FC<±1.5 were also considered in data analysis, specifically when considering overlapping gene expression groups in figure 3 (labeled as Sig<1.5FC), to include significant genes with less than ±50% alteration in expression-level that may nonetheless be biologically significant.

### Immunohistochemistry

Animals were transcardially perfused with saline and 4% paraformaldehyde (PFA, Sigma-Aldrich) in 0.1M PBS. The brain was extracted and placed in 4% PFA for 24-hrs, then dehydrated in 30% sucrose prior to sectioning. Coronal sections of the brain were obtained using a cryostat (Leica Microsystems) at 40μm slice thickness. Slices were blocked and permeabilized in PBS containing 5% normal goat serum (Cell Signaling) and 0.4% Triton-Rx100 (Alfa Aesar) for 1-hour. Primary antibodies were applied at 1:500 overnight at 4°C, Alexa-Fluor conjugated secondary antibodies (Life Technologies) were applied at 1:1000 overnight at 4°C. DAPI was applied at 1:10,000 dilution for 45-minutes and sections mounted with VectaShield HardSet (H-1400, Vector Laboratories). Antibodies included Activated Beta Catenin (05-665, Millipore Sigma), Doublecortin (AB2253, Millipore Sigma). Images were obtained by confocal microscopy (Olympus FV1000, 20x, 405nm 5%, 473nm 3%, 559 8%), all images were obtained at the same settings.

## Results

### Mesial temporal lobe epilepsy model and experimental design

In the intra-hippocampal kainate model of epileptogenesis, spontaneous recurrent seizures occur 10-14 days after SE-induction (Twele, 2016a; Twele, 2016b). We therefore hypothesized that critical gene expression changes that may be responsible for the development of epilepsy may occur within this latent period. To identify differentially expressed genes at the earliest stages of epileptogenesis, we focused our investigation on the first 2-weeks after SE-induction. Since the dentate gyrus has been specifically implicated in temporal lobe epilepsy (Parent, 1997; Kralic, 2005; Kron, 2010; Cho, 2015; Krook-Magnuson, 2015), we performed anatomical microdissection and extracted the dentate gyrus from kainate and saline-vehicle control injected animals for analysis. Time-points sampled include 3-days, 7-days, and 14-days, representing the end of SE, the period between SE and the onset of recurrent seizures, and the period near the onset of recurrent seizures respectively (Twele, 2016b). It has been shown previously that immature dentate granule cells in the contralateral dentate gyrus also undergo remodeling 2-weeks after SE and, as a result, may contribute to the maintenance of the seizure network (Wendling, 2010; Nadler, 2014; Sheybani, 2018; Gupta, 2019). Therefore, transcriptional dysregulation in both the ipsilateral dentate gyrus epileptogenic zone (EZ) and contralateral dentate gyrus peri-ictal zone (PIZ) were assessed (Figure 1A).

**Figure 1.**
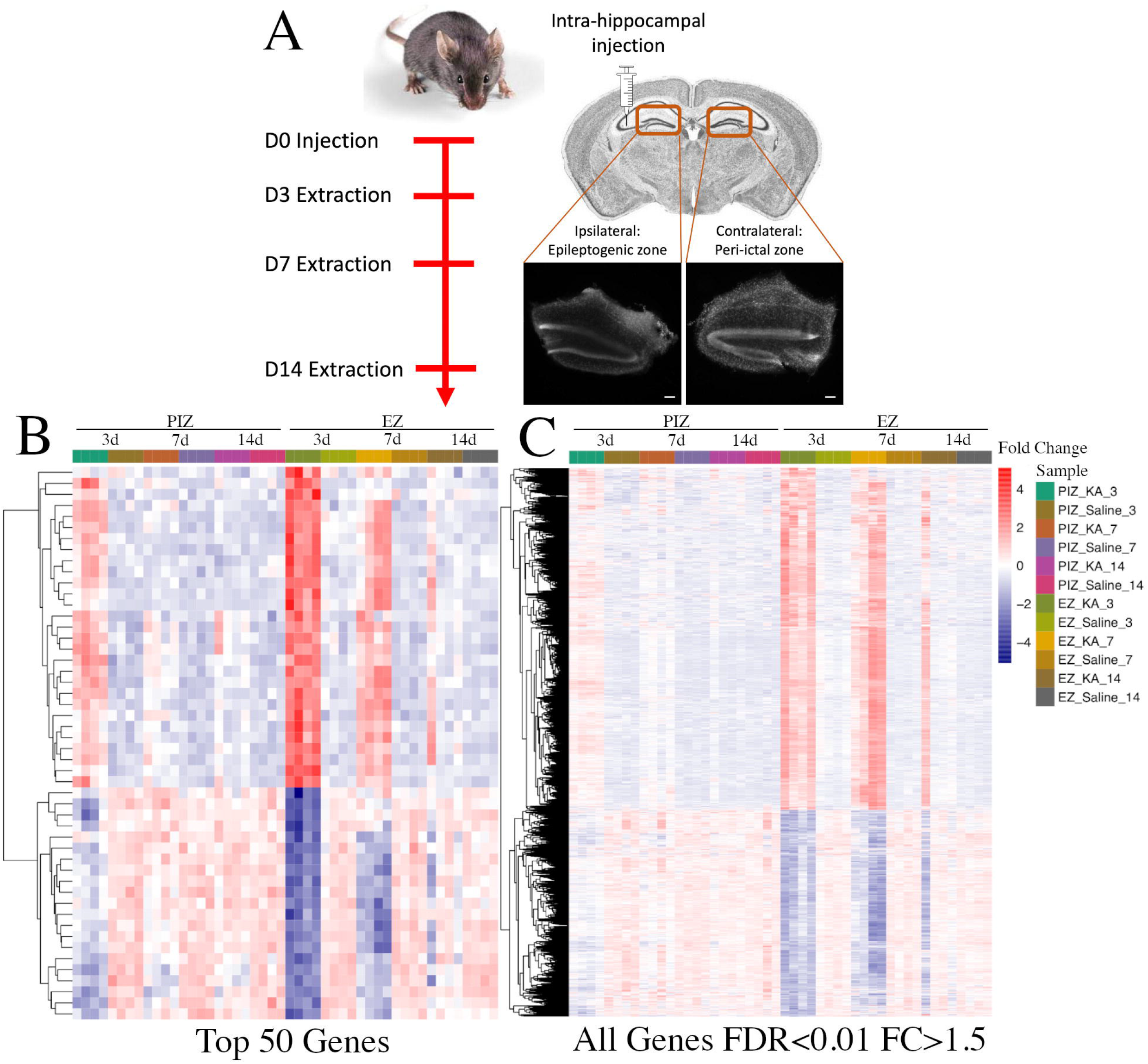
RNA-sequencing of the ipsilateral and contralateral hippocampal dentate gyri during epileptogenesis. A) Timeline and schematic of the experimental design. Mice were injected unilaterally in the hippocampus with kainate to induce SE or saline control and sacrificed 3-days, 7-days and 14-days after injection. The bilateral dentate gyri were harvested and the dorsal portions underwent RNA-sequencing, images demonstrate representative brightfield coronal sections of micro-dissected dentate gyri, scale bar 100μm. B) Hierarchical cluster analysis demonstrating B) the top 50 genes and C) genes across the entire transcriptome with FDR<0.01 and FC>±1.5 dysregulated in the EZ 3-days after SE across all samples, n=4 mice per group, 2 males and 2 females.

### Transcriptomic dysregulation across epileptic tissue regions in early epileptogenesis

We initially examined the relative expression of the most differentially expressed genes between epileptic animals and saline-injected control animals between 3 to 14-days after induction of status epilepticus (Figure 1B). In doing so, we were able to broadly assess the overall patterns of gene dysregulation over the first 2-weeks of epileptogenesis associated with both the epileptogenic zone and the peri-ictal zone. The gene expression heatmap demonstrate that in the epileptogenic zone, within the top 50 differentially expressed genes 58% are upregulated (29/50) and 42% are downregulated (21/50), 3-days after SE-induction. This pattern is maintained (though with an overall reduction in fold-change magnitude) in the epileptogenic zone 7-days after SE-induction, and begins to normalize 14-days after SE-induction. When we assess the peri-ictal zone, we observe a similar pattern of differential expression in the same subset of genes that are dysregulated in the epileptogenic zone 3-days after SE-induction, at an overall reduced fold-change magnitude. The magnitude of dysregulation of these top 50 genes in the peri-ictal zone 3-days after SE-induction is comparable to the levels observed 7-days after seizure induction in the epileptogenic zone. In the peri-ictal zone, gene expression changes begin to subside at 7-days and normalize by 14-days after SE-induction when compared to saline controls. These patterns of gene expression are also observed across the entire transcriptome (Figure 1C), and when assessed at the other sampled time points.

These data suggest that unique transcriptional dysregulation events occur within the first 2-weeks after SE-induction; these may be important in the development of spontaneous recurrent seizures, which occurs toward the end of this latent period. These data also suggest that some transcriptional changes are shared between the epileptogenic zone and seizure network, and that transcriptional dysregulation is of greater magnitude and prolonged in the epileptogenic zone.

### Enrichment of functional DAVID categories in the epileptogenic zone and seizure network across early epileptogenesis

We next examined the enrichment of gene ontology (GO) terms within the DAVID pathways, in order to investigate the functional effect of gene dysregulation at each timepoint across the two regions. We specifically identified the top 50 most significantly enriched GO categories within each of the 2 regions (Figure 2A, 2B), and also compared the pattern of expression across the 2 regions using the top 50 enriched categories in the epileptogenic zone as reference (Figure 2C). We observe that in the epileptogenic zone, different overall patterns of GO terms are shared between the 3-day and 7-day timepoints, and between the 7-day and 14-day timepoints, with a number of GO terms unique to the 14-day time-point. In the epileptogenic zone, GO terms enriched at 3-days and 7-days after SE are associated with cell signaling (e.g. intracellular signal transduction, regulation of signaling, regulation of cell communication, regulation of signal transduction, positive regulation of molecular function, positive regulation of signal transduction, regulation of intracellular signal transduction), cell migration and motility (e.g. cell migration, cell adhesion, movement of cell or subcellular component, regulation of cell motility, regulation of locomotion, locomotion, cell motility, localization of cell), as well as cell survival/proliferation (e.g. cell proliferation, neurogenesis, cell death, apoptotic process, programmed cell death, cell development) (Figure 2A). These GO terms encompass pathological cellular processes observed in the dentate gyrus after SE (Buckmaster, 1997), and suggest that the molecular mechanisms responsible for these pro-epileptogenic processes may be regulated at these early time-points after SE. GO terms enriched at the 7-day and 14-day timepoints, as well as more prominently in the 14-day timepoint, are related to inflammatory processes (e.g. immune response, response to cytokine, inflammatory response, regulation of TNF superfamily cytokine production, regulation of adaptive immune response and more). These inflammatory GO terms are less represented at the 3-day timepoint, and if present, overlap with the 7-day and 14-day timepoints. This suggests that pro-inflammatory processes are initiated in a delayed and prolonged manner from 7-days after SE in the epileptogenic zone.

**Figure 2.**
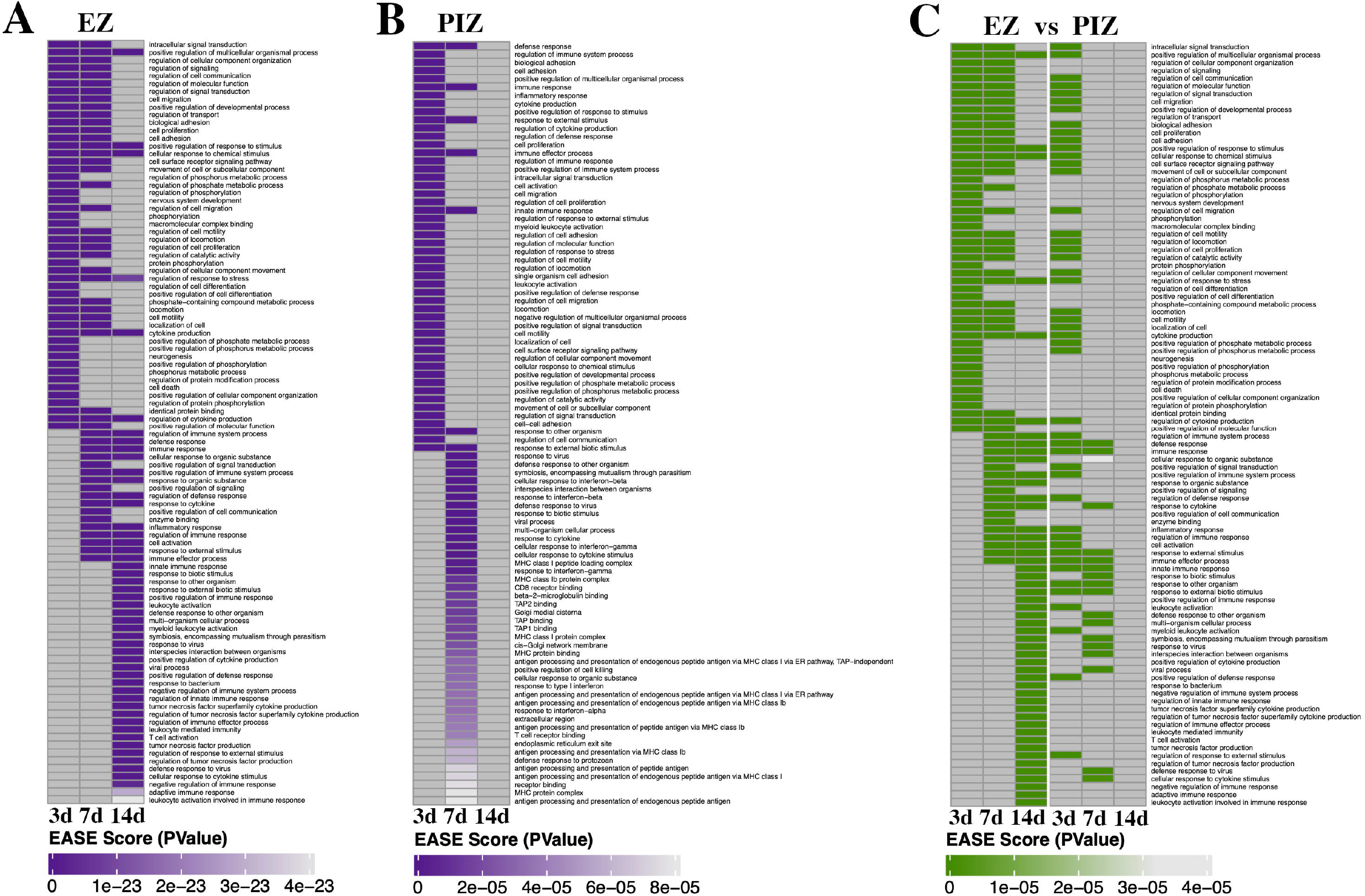
Gene-set enrichment in the EZ and PIZ over time. Significant DAVID terms are expressed over the first 2-weeks of epileptogenesis as heatmaps across the EZ and PIZ. A) Enrichment of gene-sets in the EZ, 3-days 7-days and 14-days after SE-induction. B) Enrichment of gene-sets in the PIZ, 3-days, 7-days and 14-days after SE induction. C) Corresponding PIZ gene-set fold-change in the most-highly enriched EZ gene-sets.

**Figure 3.**
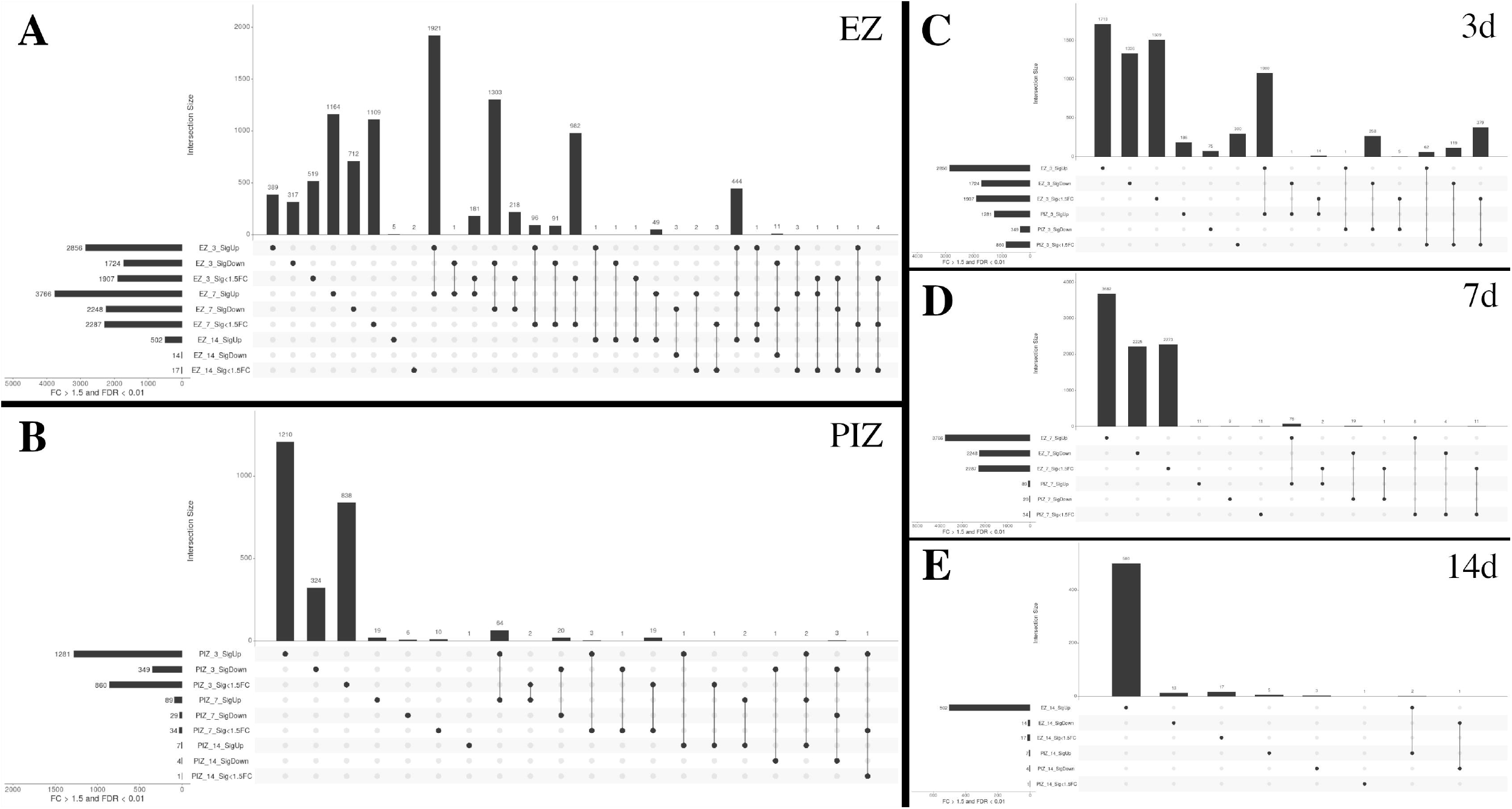
UpSet plots depicting the extent of shared transcriptional dysregulation in the EZ and PIZ across the first 2-weeks of epileptogenesis. For each UpSet plot group, seizure network region (EZ/PIZ), time point in days (3, 7, 14) and differential expression group (SigUp, SigDown, Sig<1.5FC) are labeled. The dots denote genes specific to each individual group, linked dots represent overlapping gene groups. The vertical bars demonstrate gene number for the corresponding individual or linked groups. The horizonal bars denote total gene number for each region/time-point/differential expression group. Transcriptional dysregulation across A) the EZ and B) the PIZ, 3-days, 7-days and 14-days after SE-induction. Transcriptional dysregulation across overlapping and unique groups between the EZ and PIZ C) 3-days, D) 7-days, E) 14-days, after SE-induction

Within the peri-ictal zone, the majority of the top 50 significantly enriched GO terms are represented uniquely at 3-days or 7-days after SE-induction with few overlapping groups, and none represented at 14-days (Figure 2B). These data further support the return of transcriptional dysregulation to baseline 14-days after SE-induction. The GO terms enriched 3-days after SE-induction are, similarly to the 3-day/7-day timepoint in the ipsilateral region, associated with cell signaling (e.g. intracellular signal transduction, positive regulation of signal transduction, cell surface receptor signaling pathway, regulation of signal transduction, regulation of cell communication, positive regulation of cell signaling), cell migration and motility (e.g. cell adhesion, cell migration, regulation of cell motility, regulation of locomotion, positive regulation of cell adhesion) and cell survival/proliferation though with less representation of cell death related GO-terms as might be expected in comparison the ipsilateral epileptogenic zone (e.g. cell proliferation, regulation of cell proliferation, positive regulation of developmental process). There is, however, earlier representation of immune/inflammation related GO terms (e.g. regulation of immune system process, immune response, cytokine production, leukocyte activation, regulation of inflammatory response) 3-days after SE-induction, which also comprise the predominantly represented terms 7-days after SE in the peri-ictal zone (e.g. response to cytokine, cellular response to interferon gamma, antigen processing and presentation of peptide antigen).

When we examine how the most highly enriched GO-terms in the epileptogenic zone are expressed in the PIZ across the 3, 7 and 14-day timepoints (Figure 2C), we observe that there are similar functional consequences between the ipsilateral and contralateral regions 3-days after SE-induction. We observe enrichment of GO-terms focused on cell signaling (e.g. intracellular signaling induction, regulation of signaling, regulation of cell communication, regulation of molecular function, regulation of signal transduction) and cell migration and motility (e.g. cell migration, biological adhesion, regulation of cell migration, regulation of cell motility, regulation of locomotion), most of which are also represented 7-days after SE-induction in the epileptogenic zone (Figure 2C). These data suggest that early cell signaling processes may be critical for the initiation of pro-epileptogenic remodeling in both the epileptogenic zone and peri-ictal zone. Days 3 & 7 in the peri-ictal zone and days 7 & 14 in the epileptogenic zone share overlapping enrichment of immune/inflammation related GO terms (e.g. immune response, response to cytokine, immune effector process, cellular response to cytokine stimulus). Our data therefore suggest that inflammation may also play an early role in epileptogenesis, with expression of inflammation related gene-sets occurring in both the epileptogenic zone and peri-ictal zone within the first 14 days after SE-induction.

### Shared patterns of gene dysregulation between the epileptogenic zone and peri-ictal zone over early epileptogenesis

We examined overlapping gene sets to determine whether differentially expressed genes were shared across time and anatomical region, potentially indicative of shared underlying molecular processes. In the data represented in figure 3, significantly upregulated genes are labeled as “SigUp”; significantly downregulated genes are labeled “SigDown”, and genes with FDR<0.01 but with less than 50% fold change are labeled as “Sig<1.5FC”. Over early epileptogenesis in the epileptogenic zone, the largest overlapping groups were observed between 3-days and 7-days after SE-induction. This indicates a large degree of overlap in underlying gene dysregulation, and therefore likely shared molecular processes, in the first week after SE in the epileptogenic zone. The number of dysregulated genes that remain either upregulated or downregulated 14-days after SE-induction falls by over 85%, suggesting that the critical period after SE may be captured within the first week (Figure 3A). By comparison, in the peri-ictal zone the majority of genes are uniquely dysregulated 3d after SE-induction, with few dysregulated genes observed at all other time-points; we observe less overlap between the 3d and 7d timepoints, with overlap near-negated at the 14d timepoint (Figure 3B). These data suggest that the principal gene dysregulation period in the peri-ictal zone may occur within 3d of SE-induction.

When compared across region at each timepoint, the epileptogenic zone and peri-ictal zone demonstrate differing patterns of gene dysregulation. Three-days after SE-induction, there were 1348 overlapping genes between significantly upregulated and downregulated genes in the epileptogenic zone and peri-ictal zone (Figure 3C). This suggests that some gene dysregulation events and their corresponding pathways are shared between these two regions of the seizure network 3-days after SE-induction. However, the number of overlapping genes reduced 7-days (Figure 3D) and 14-days (Figure 3E) after SE-induction, suggesting that gene dysregulatory events diverge beyond 3-days after SE. Fourteen days after SE, 502 genes were upregulated in the epileptogenic zone, 500 uniquely and only 2 were shared with the peri-ictal zone (Figure 3E); however, 444 genes overlapped with earlier time-points in the epileptogenic zone 3-days and 7-days after SE (Figure 3A). These data indicate that transcriptional dysregulation in the peri-ictal zone declines 14-days after SE-induction, and the epileptogenic zone retains persistent gene dysregulation, as might be expected from the extent of injury and remodeling in the epileptogenic zone 2-weeks after SE-induction.

### Wnt pathway dysregulation during early epileptogenesis

Previous work has implicated the Wnt pathway as a factor in neuronal remodeling in the dentate gyrus early after SE-induction in this model (Gupta, 2019). Furthermore, Wnt pathway gene changes have also been identified in adult and infantile generalized epilepsy models (Theilhaber, 2013; Dingledine, 2017; Canto, 2021), suggesting that Wnt pathway gene changes may be relevant in adult focal epilepsy. We therefore specifically examined the transcriptional dysregulation of Wnt pathway genes across the epileptogenic zone and peri-ictal zone.

The modified KEGG pathway presented demonstrates gene expression data from the epileptogenic zone (Figure 4A) and peri-ictal zone (Figure 4B) 3-days after SE-induction. Positive upregulation of canonical and non-canonical Wnt signaling mediators and downstream effector genes is observed in both the epileptogenic zone and peri-ictal zone 3-days after SE (Figure 4A, 4B). In the epileptogenic zone, canonical pathway positive effector genes such as Fzd1 (FC 3.09, p-adjusted 1.92×10^-7), LRP5 (FC 2.03, p-adjusted 5.17×10^-4), Lef1 (FC 1.63, p-adjusted 9.29×10^-3) and CTNNB1/beta-catenin (FC 1.24, p-adjusted 9.80×10^-4) are noted to be upregulated and negative regulator genes such as GSK3b (FC −1.47, p-adjusted 1.19×10^-5), Btrc (encoding beta-TrCP (Marikawa, 1998; Liu, 1999)) (FC −1.39, p-adjusted 2.30×10^-4), Wif1 (FC −4.59, p-adjusted 3.50×10^-3), DKK3 (FC −2.70, p-adjusted 6.12×10^-11) and APC (FC −1.37, p-adjusted 7.48×10^-3) are downregulated, suggesting that the canonical Wnt pathway may be activated 3-days after SE in the epileptogenic zone. Non-canonical Wnt pathway genes are also dysregulated with some genes upregulated, such as Wnt5b (FC 2.09, p-adjusted 3.35×10^-2), ROR2 (FC 4.47, p-adjusted 1.35×10^-2), RhoA (FC 1.67, p-adjusted 6.13×10^-7), Prickle3 (FC 2.18, p-adjusted 3.49×10^-2), and other genes downregulated, such as Prickle2 (FC −1.85, p-adjusted 3.02×10^-6), JNK (FC −1.69, p-adjusted 3.46×10^-6), PKC (FC – 2.42, p-adjusted 6.36×10^-11) and CaMKII (FC −2.80, p-adjusted 4.53×10^-7).

**Figure 4.**
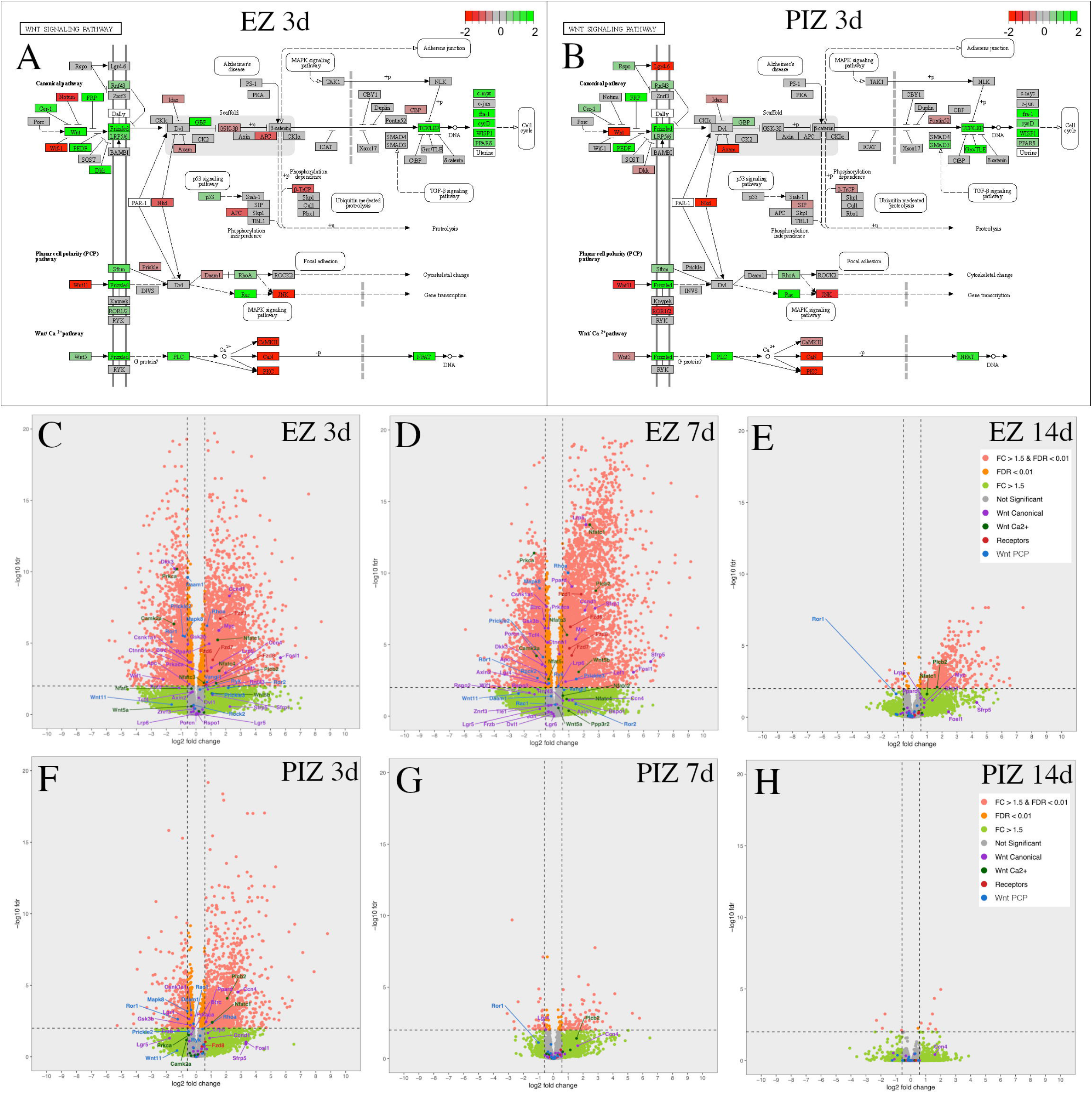
Wnt pathway transcriptional dysregulation in the EZ and PIZ during the first 2-weeks of epileptogenesis. Modified KEGG pathways representing canonical and non-canonical planar cell polarity and calcium pathway signaling 3-days after SE-induction in the A) EZ and B) PIZ. Scale bar represents log2-fold-change in transcription, green color denotes upregulation and red color denotes downregulation. Whole transcriptome volcano plots with Wnt signaling mediators individually labeled in the EZ C) 3-days, D) 7-days and E) 14-days, and in the PIZ F) 3-days, G) 7-days, and H) 14-days after SE-induction, dashed lines denote FDR<0.01 & FC>±1.5 (equivalent to log2-fold-change>±0.585). Pink dots represent genes with FDR<0.01, FC>±1.5; orange dots represent genes with FDR>0.01, FC<±1.5; green dots represent genes with FDR>0.01, FC>±1.5; gray dots represent genes with FDR>0.01, FC<±1.5.

Transcriptional dysregulation of these Wnt pathway genes was followed across the epileptogenic zone (Figure 4C-E) and peri-ictal zone (Figure 4F-H) at each time-point 3-days, 7-days and 14-days after SE (Figure 4B-G). Canonical Wnt gene dysregulation is most highly represented in the epileptogenic zone 3-days and 7-days after SE (Figure 4A, 4C, 4D). Non-canonical genes are also dysregulated in the epileptogenic zone 3-days and 7-days after SE, with fewer genes significantly altered in expression (FC>±1.5, FDR<0.01) than in the canonical Wnt pathway (Figure 4A, 4C, 4D). Frizzled receptor genes are upregulated in the epileptogenic zone 3-days and 7-days after SE (Figure 4C, D). In the peri-ictal zone, fewer canonical and non-canonical Wnt genes are dysregulated 3-days after SE than in the epileptogenic zone; none are represented 7-days after SE (Figure 4F, G). Wnt pathway genes are not significantly dysregulated in either region 14-days after SE (Figure 4E, H). As Wnt gene dysregulation appears to reduce progressively over the first 14-days after SE, these data suggest that the first 7-days after SE may represent the therapeutic window for potential Wnt-modulation based interventions.

### Canonical Wnt signaling is upregulated early after SE by expression of activated-beta catenin

Our transcriptomic analysis suggests that canonical Wnt pathway expression is increased 3-days and 7-days after SE-induction in the epileptogenic zone, and to a lesser extent in the peri-ictal zone 3-days after SE. In order to determine whether the transcriptional changes we observe in the Wnt pathway reflect changes in molecular signaling during epileptogenesis, we evaluated the expression of the active form of beta-catenin, the central signaling molecule in the canonical Wnt pathway (Oliva, 2013). We evaluated activated beta-catenin through immunohistochemistry in the dentate gyrus 3-days, 7-days and 14-days after SE-induction (Figure 5). As expected, granule cell dispersion was evident in the epileptogenic zone 7 and 14-days after SE, with loss of DCX-positive dentate granule cells (Figure 5C, 5D) (Bouilleret, 1999; Jessberger, 2005; Kralic, 2005). We observe expression of active beta-catenin 3-days after SE-induction both in the peri-ictal zone (Figure 5F) and, more strongly, in the epileptogenic zone, especially in the hilus (Figure 5B). Interestingly, active beta-catenin expression diminishes at later time-points (Figure 5C, 5D, 5G, 5H), consistent with our KEGG-pathway data (Figure 4A, 4B) and individual geneexpression transcriptional data. These results validate the finding of early canonical Wnt activation as identified in the transcriptomic data.

**Figure 5.**
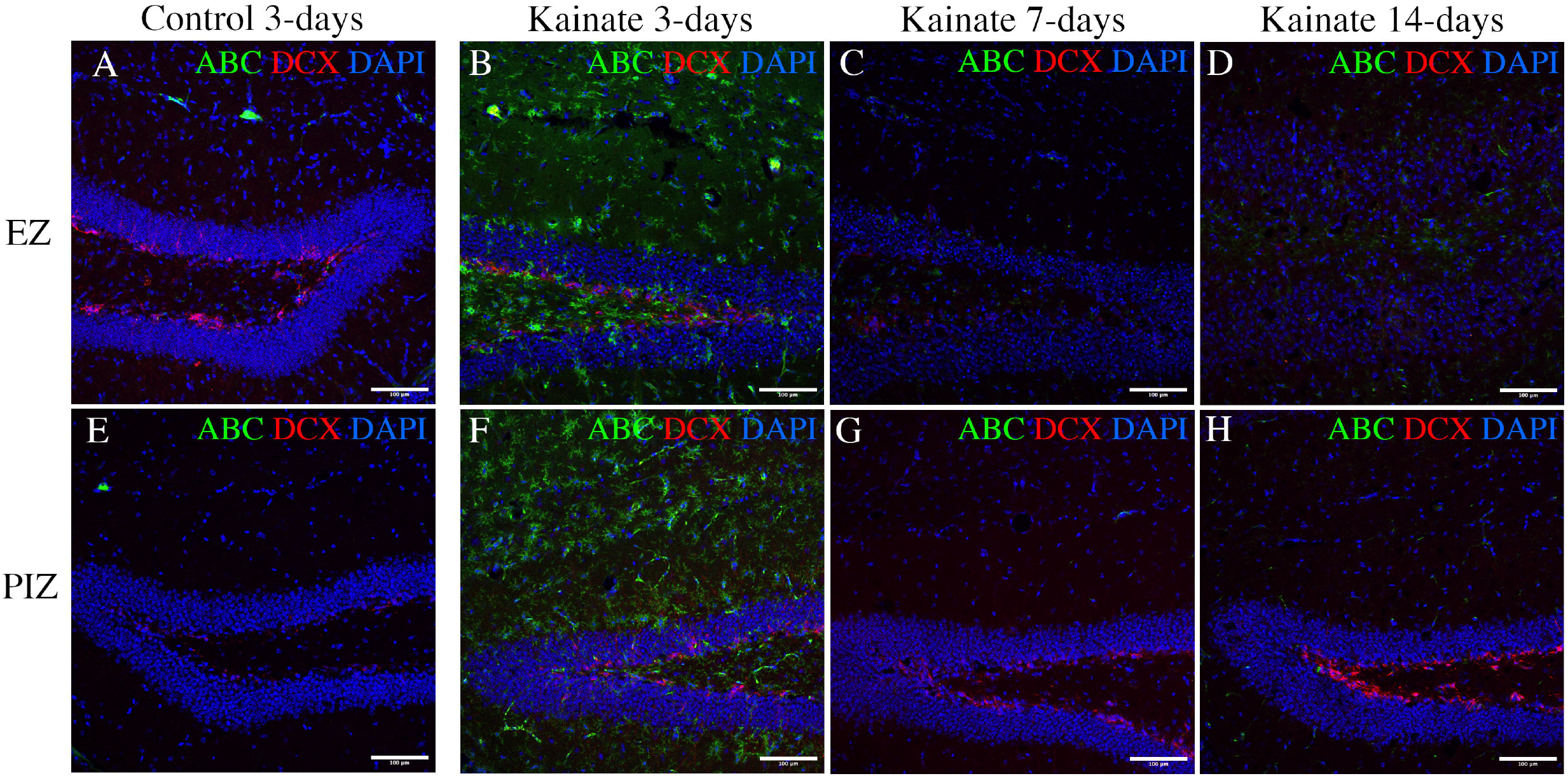
Immunohistochemical evidence for canonical Wnt activation in the hippocampus. Immunofluorescent images demonstrating active-beta-catenin (ABC – a marker of canonical Wnt pathway activation) and doublecortin (DCX – a marker of immature dentate granule cells) in A-D) the EZ (top row), and E-H) PIZ (lower row) dentate gyrus. Images demonstrate active beta-catenin expression (green) A, E) in control mice 3-days after saline injection (7-days and 14-days unchanged, data not shown), and in kainate-injected mice B, F) 3-days, C, G) 7-days and D, H) 14-days after SE. Scale bar 100μm.

## Discussion

There is increasing interest in understanding the molecular pathways underpinning pathological remodeling in epileptogenesis. Mechanistic insights into the pathways responsible will be critical for understanding the disease and generating novel therapeutic strategies to prevent the development of epilepsy. Such therapeutic approaches have the potential to be extremely valuable in the setting of medical conditions at high risk of developing recurrent seizures, for example after presentation with first seizure, and other neurological conditions such as febrile seizure, infection, tumor and trauma (Frey, 2003; Myint, 2006; French, 2012). Furthermore, established clinical treatments for epilepsy, such as medications and surgery can be ineffective and can have adverse neuropsychological consequences (Helmstaedter, 2004; Drane, 2015).

In this study we focused our investigation on the pre-clinical intrahippocampal kainate mouse model of epilepsy, as a model for clinical focal temporal lobe epilepsy. It has been shown previously that the hippocampus injected with kainate undergoes extensive pathological remodeling of the dentate gyrus consistent with epileptogenic foci in focal temporal lobe epilepsy, and the contralateral hippocampus undergoes pathological remodeling events consistent with seizure network changes (Gupta, 2019) and can generate independent seizures (Häussler, 2012), features seen and targeted in clinical temporal lobe epilepsy (Wendling, 2010; Sheybani, 2018). Like other studies (Li, 2010; Dingledine, 2017; Canto, 2021), we isolated the dentate gyrus as it undergoes extensive remodeling in epileptogenesis and is believed to be responsible for spontaneous recurrent seizures (Buckmaster, 1997; Parent, 1997; Choi, 2005; Kralic, 2005; Cho, 2015). Uniquely, our study design allows investigation of different regions of the seizure network, including the bilateral dentate gyri representing the epileptogenic zone and peri-ictal zone, providing insights into their relative contributions to epileptogenesis.

Our data interestingly demonstrate shared patterns of transcriptomic dysregulation in the epileptogenic zone and the peri-ictal zone during the first week of epileptogenesis. Bilateral neuronal activation in the dentate gyrus has previously been demonstrated within hours of status epilepticus induction in this model, and these gene changes may well represent activation-dependent changes. These early changes, however, <u>decline</u> in both the epileptogenic zone (EZ) and peri-ictal zone (PIZ) by 2-weeks after SE, which is sufficient time for extensive remodeling to be observed in the epileptogenic zone and peri-ictal zone (Gupta, 2019), suggesting that these early transcriptional changes may contribute to epileptogenic remodeling.

The patterns of transcriptional dysregulation shared between the epileptogenic zone and peri-ictal zone include enrichment of cell signaling, motility and proliferation functional categories, as might be expected given the pathological remodeling of dentate granule cell neurons observed in the dentate gyrus during epileptogenesis. Interestingly these processes decline within 14-days of SE-induction, suggesting that the window to modulate these processes therapeutically may be brief. These data also have implications for the use of the contralateral hippocampus as a control. We observe early transcriptional changes in the peri-ictal zone that correlate with those that occur within the epileptogenic zone, which return to baseline within 7-14-days. Therefore, sampling outside of this period may not capture these early gene changes. As it has been demonstrated previously that the contralateral hippocampus undergoes pathological remodeling and generates seizures after ipsilateral kainate injection (Häussler, 2012; Gupta, 2019; Berger, 2020), it is possible that these gene changes may have delayed functional consequences. Immune processes followed a different time-course within the epileptogenic zone and peri-ictal zone, with delayed representation in the epileptogenic zone beginning 7-days after SE sustained to 14-days after SE; whereas in the peri-ictal zone, immune processes were represented 3-days and 7-days after SE, and no longer represented 14-days after SE. These findings indicate that inflammation may also play a role in altering neuronal phenotype and warrant further investigation.

Previous work has investigated the role of the Wnt pathway in epileptogenesis, utilizing this model, and demonstrated that specific Wnt pathway ligands are differentially expressed in the epileptogenic zone and peri-ictal seizure network 3-days after SE-induction (Gupta, 2019). Data from our current study corroborate these published findings with upregulation of Wnt5b in the epileptogenic zone, Wnt7b upregulation across both the epileptogenic zone and peri-ictal zone, and Wnt9a downregulation across both the epileptogenic zone and peri-ictal zone, providing evidence for the robustness of the model. The whole transcriptome data in this study suggest that canonical Wnt signaling mediators are upregulated in the epileptogenic zone during epileptogenesis, either in a causative capacity or reactive to SE. We observed expression of active beta-catenin protein by immunofluorescence in the epileptogenic zone and peri-ictal zone 3-days after SE-induction; this correlated with gene expression data for beta-catenin in the epileptogenic zone, but not in the peri-ictal zone. Beta-catenin is regulated both by expression but also by phosphorylation status and degradation by GSK3b (Oliva, 2013); therefore, transcription level of beta-catenin alone may not fully reflect signaling activity and activity of downstream genes. Direct assessment for active beta-catenin can provide a more accurate picture, as demonstrated in our immunohistochemistry and KEGG pathway data. The specific role of Wnt pathway dysregulation in epileptogenesis remains to be fully elucidated; given the complexity of the signaling pathways, including mixed changes in expression of the non-canonical pathways and inhibitor molecules evidenced in our data, it is possible that the role of Wnt pathway dysregulation in epileptogenesis may be temporally defined, as it is during hippocampal neurogenesis (Arredondo, 2020a). Furthermore, different Wnt pathways and signaling mediators may play different roles in the formation of the epileptogenic zone and peri-ictal zone. Indeed, Wnt signaling has been reported as both protective and deleterious in rat generalized epilepsy models; network/region specific changes remain to be fully elucidated (Fasen, 2002; Madsen, 2003; Sun, 2020; Alqurashi, 2021; Jean, 2022).

Several studies on RNA sequencing in animal pre-clinical epilepsy models have contributed to our understanding of gene expression changes at various stages of epilepsy. Canto et al., 2021, investigated transcriptomic and proteomic changes 15-days after SE, during the epileptogenic period, using the rat systemic pilocarpine generalized temporal lobe epilepsy model; tissues were separated into dorsal and ventral dentate gyrus and CA3 sub-regions. The investigators observed a decrease in neuronal markers and increased microglial markers in the DG and CA3 sub-regions, and specifically implicated NMDA excitotoxicity, calcium/calmodulin dependent protein kinase (CaMK) and LRRK2/Wnt signaling. Interestingly, this study reported increased expression of CaMKs and LRRK2, which was observed in our data, as well as specific enrichment of the pathway “Adult neurogenesis in the subventricular zone” in the dentate gyrus, which they concluded to be mediated by LRRK2, Wnt and CDK5. LRRK2 is a Wnt signaling scaffold protein that has been implicated in the pathogenesis of Parkinson’s disease (Salasova, 2017), and mediates canonical Wnt signaling (Berwick, 2012). Other Wnt-related processes were also upregulated in the DG, including NFKB and TGF-beta signaling.

Given the extensive variability in pre-clinical epilepsy models, Dingledine et al., 2017, utilized a number of rat pre-clinical generalized epilepsy models, including systemic pilocarpine, kainate and electrical kindling models, and extracted the mid-layer of the right dentate gyrus for RNA-sequencing. Of the 72 dysregulated genes common to all models, 19 genes were related to Wnt signaling. These genes include ABCD2 (Park, 2013), C1s (Naito, 2012), CBFB (Xia, 2018), CRIM1 (Ponferrada, 2012), Ext1 (Wang, 2019), FAM129B (Conrad, 2013), FAT1 (Morris, 2013), GDF10 (Katoh, 2006), GPC3 (Gao, 2011), KHDRBS3 (Ukai, 2021), KLF15 (Noack, 2012), Lox (Che, 2018), MMP9 (Ingraham, 2011), NTF3 (Patapoutian, 1999), SS18 (Cironi, 2016), SSBP3 (van Tienen, 2017), TRH (Skah, 2017), Wls (Das, 2012), ZMIZ1 (Lomeli, 2022). Of these genes, 17 are represented in our transcriptomic data, highlighting the potential overlap in molecular signaling identified in their study and ours. They did not, however, specifically implicate the Wnt pathway. Their methods investigated the mid-layer of the dentate granule cell layer by laser microdissection. While specific for the dentate, this method excluded the region adjacent to the hilus containing immature dentate granule cells, which are known to undergo remodeling during epileptogenesis (Overstreet-Wadiche, 2006; Kron, 2010; Gupta, 2019). Our histological examination demonstrates that, within the dentate gyrus, active-beta-catenin (marker of canonical Wnt signaling activation) is predominantly expressed in this region adjacent to the subgranular zone and less so in the mid-dentate gyrus. This provides some suggestion as to why Wnt signaling may have been less represented in their study.

Hansen et al., 2014 performed RNA sequencing of the whole hippocampus using the systemic pilocarpine mouse model of generalized temporal lobe epilepsy. In this study, the investigators sampled the whole hippocampus, unlike the previous two studies which focused on the dentate gyrus. Nevertheless, the authors of this study acknowledged the central role of the dentate gyrus in epileptogenesis. Timepoints were selected across various periods of epilepsy, 12hrs representing acute SE, 10-days representing the latent period and 6-wks representing chronic epilepsy. As might be expected, the investigators found that the 3 time-points were relatively different in transcriptomic profile, likely as they represent different phases of pre-clinical model epilepsy development. Interestingly, they found increased similarity between the 12-hr acute SE and 6-wk chronic phase, which the authors conclude maybe due to ictal activation related changes; the 10d time-point representing a unique phase specific to epileptogenesis. These insights and others guided our choice of timepoints within the latent epileptogenic period (Okamoto, 2010; Hansen, 2014; Twele, 2016b). In addition to the previously discussed studies (Dingledine, 2017; Canto, 2021), work by Theilhaber et al., 2013 also implicated Wnt signaling in pre-clinical epileptogenesis. Using a neonatal rat pup hypoxia model of generalized epilepsy, the whole hippocampus and cortex were analyzed by RNA-sequencing; the investigators observed increased expression of canonical Wnt signaling molecule genes such as beta-catenin, LRP6, Dvl3, as well as inhibitors such as GSK3b, sFRP2 and Wif1. Notably in this model, hippocampal cell death is not observed after hypoxic seizure, which differs from kainate-based models (Sanchez, 2001; Koh, 2004). Nonetheless, the findings of Wnt dysregulation across different pre-clinical epilepsy models potentially suggest that the Wnt pathway may be relevant in the pathogenesis of epilepsy.

Our use of a focal pre-clinical model of mTLE additionally allows investigation of different regions of the seizure network, including the epileptogenic zone and peri-ictal zone. This may enhance clinical translation, as there is increasing awareness of the role of the seizure network in human clinical epilepsy, with network nodes being targeted therapeutically and being held responsible for seizure recurrence after epilepsy surgery (McIntosh, 2004; Wendling, 2010; Barba, 2016; Bjellvi, 2019). Our data demonstrate that there are both similarities and differences between different sub-regions of the seizure network, suggesting that epileptogenic processes may follow common pathways. The canonical Wnt pathway is implicated in the formation of the seizure network, whether in a causative or reactive manner remains to be determined in future studies.

In conclusion, our study investigates transcriptomic changes across a focused period during epileptogenesis, the latent period between status epilepticus and the development of spontaneous recurrent seizures. We differentiate between different regions within the seizure network, utilizing the intrahippocampal kainate model for focal mesial temporal lobe epilepsy to examine the epileptogenic zone and the peri-ictal zone. We specifically examined Wnt pathway dysregulation and found canonical Wnt pathway activation in the epileptogenic zone and peri-ictal zone early in the latent period. Many questions remain, is there re-emergence of various signaling pathways during breakthrough clinical seizures or during recurrence after epilepsy surgery, can these pathways be modulated to alter the course of epileptogenesis, do various clinical presentations and pre-clinical models of epilepsy share common signaling pathways. Our database can be interrogated to determine the involvement of other molecular pathways in the formation of the seizure network. There remains a need for improved graphic representation, data integration amongst studies and defined epilepsy-specific gene-sets for data analysis interpretation. With these, a more integrated view of epileptogenesis may be achieved.

## Limitations

There are limitations of our study that ought to be considered in interpretation of the data. The pre-clinical focus uniquely allows access to mammalian hippocampal tissues at early timepoints during epileptogenesis, which cannot be routinely accessed in human epilepsy patients as time to presentation for epilepsy surgery can be 20-years from first presentation (Wiebe, 2001). Variation in pre-clinical models and their concordance with clinical epilepsy must therefore be considered; the presence of pathological changes in the dentate gyrus of epileptic mice and humans is now well-documented with shared histological features such as granule cell dispersion, hippocampal sclerosis, aberrant migration and dendrite formation (Houser, 1990; Parent, 1997; Mathern, 2002; Crespel, 2005; Blumcke, 2007; Kron, 2010; Blumcke, 2013; Cho, 2015). These lend biological credibility to pre-clinical animal models; however, human studies are still needed to validate pre-clinical findings and generate hypotheses for development of novel therapeutics targeting epileptogenesis. Human tissue studies have been reported, and though interpretation of findings is limited by the use of cadaveric and allogenous tumoral tissues as controls, Wnt pathway genes such as casein kinase have been observed in human mTLE (Dixit, 2016; Vangoor, 2019). Attempting to increase the fidelity of pre-clinical models to human clinical epilepsy is therefore of value to improve translation. Our utilization of a focal model of mesial temporal lobe epilepsy as a model for human mTLE aims to address this; however, the use of kainate injection may not accurately recapitulate the typical etiology in humans. Nonetheless, insights can be made into the epileptogenic process.

Given the role of the dentate gyrus in epileptogenesis, our study focused on microdissected dentate gyrus tissue as described previously (Gupta, 2019), as opposed to laser microdissection studies that focus on specific sub-regions of the dentate gyrus (Dingledine, 2017) and largely excluded immature dentate granule cells, the principal cell-type aberrant in pre-clinical TLE (Overstreet-Wadiche, 2006; Kron, 2010). While the dissection tissue does include the hilus region, the whole dentate is sampled and thereby includes immature and mature neurons as well as associated non-neuronal cell types, thereby reflecting the overall signaling *milieu* of the dentate gyrus. Another limitation of our study, therefore, is that in using bulk RNAseq for transcriptomic analysis, we cannot uncover cell-type specific contributions. The specific contributions of individual cell types to the signaling environment will require additional cell-specific and region-specific studies.

